# The nitrification inhibitor DMPP preferentially suppresses ammonia-oxidising bacteria and reduces nitric oxide emissions in an archaeal-dominated grassland soil

**DOI:** 10.64898/2026.09.20.753006

**Authors:** Abdallah Awad, Laura M. Cardenas, Nadine Loick, Salim Al-Babili, Mark Tester, Vanessa J. Melino

## Abstract

**Aims:** Although ammonia-oxidising archaea (AOA) dominate nitrifier communities in many agricultural soils, the contribution of the less abundant ammonia-oxidising bacteria (AOB) to nitrogen oxide emissions, particularly nitric oxide, remains poorly understood. This study investigated the roles of AOA and AOB in regulating nitric oxide (NO) and nitrous oxide (N_2_O) emissions from a urea-fertilised grassland soil and evaluated the nitrification inhibitors dicyandiamide (DCD) and 3,4-dimethylpyrazole phosphate (DMPP).

**Methods:** A 28-day soil incubation experiment was conducted using a denitrification incubation system to continuously monitor NO and N_2_O emissions. In parallel, a destructive incubation experiment was performed to determine the dynamics of ammonium (NH□□-N) and total oxidised nitrogen (TOxN) and to assess changes in the abundance of AOA and AOB following urea application with or without DCD or DMPP.

**Results:** AOA accounted for approximately 66% of the total ammonia-oxidising community. Neither inhibitor significantly affected AOA abundance, whereas both suppressed AOB, with DMPP showing stronger and more persistent inhibition than DCD. DMPP reduced NO emissions by 16–75%, whereas N_2_O emissions decreased by only 0.2– 27%. The greater reduction in NO, together with the preferential suppression of AOB, suggests that low-abundance AOB contributed disproportionately to NO production despite the numerical dominance of AOA.

**Conclusions:** These findings suggest that AOB can contribute disproportionately to NO emissions despite their lower abundance relative to AOA. The preferential suppression of AOB by DMPP was associated with a substantial reduction in NO emissions, highlighting AOB-associated processes as a potential target for mitigating reactive N losses from fertilised soils.

## 1. Background and Aims

Agricultural soils are a major source of nitrogen (N) emissions, primarily in the form of nitrous oxide (N_2_O) and nitric oxide (Cavanaugh et al.), both of which have significant environmental and agronomic impacts (Galloway et al. 2008; Ravishankara et al. 2009). Nitrous oxide is a potent greenhouse gas with a global warming potential approximately 300 times greater than carbon dioxide (Cavanaugh et al. 2022; Griffis et al. 2017). Nitric oxide plays a central role in atmospheric chemistry, as it is rapidly oxidised to nitrogen dioxide (NO_2_), contributing to tropospheric ozone formation, acid deposition, reduced plant productivity, and negative impacts on human respiratory health (Medinets et al. 2015; Perring et al. 2025; Ravishankara et al. 2009; Schreiber et al. 2012). Beyond environmental impacts, gaseous N losses reduce nitrogen use efficiency (NUE), thereby increasing fertiliser demand with economic and environmental costs(Zhang et al. 2015). Improving our understanding of the microbial processes responsible for these emissions is therefore essential for optimising N management in agricultural systems. Among these processes, nitrification, the aerobic oxidation of ammonium (NH_4_□) to nitrite (NO_2_□), is a major pathway of NO production and a significant source of N_2_O in soils(Ruser and Schulz 2015). This process is mediated by ammonia-oxidising archaea (AOA) and ammonia-oxidising bacteria (AOB) (Könneke et al. 2005; Prosser and Nicol 2012). Despite well-documented niche partitioning between AOA and AOB along gradients of NH□□ availability and soil pH, the functional consequences of this partitioning for nitrogen oxide emissions under fertilisation remain poorly resolved (Di and Cameron 2016; Leininger et al. 2006; Prosser and Nicol 2012). Mechanistic differences between AOA and AOB are well established: AOB generate NO as an obligate intermediate of hydroxylamine (NH_2_OH) oxidation and via nitrifier denitrification, whereas AOA lack the enzymatic machinery for the latter pathway and instead produce NO predominantly through abiotic reactions involving hydroxylamine and NO_2_-(Caranto and Lancaster 2017; Wrage et al. 2001). These biochemical distinctions predict substantially greater NO yields per unit of ammonia oxidised in AOB relative to AOA. N_2_O emissions are inherently more complex, arising from AOA- and AOB-mediated nitrification, including nitrifier denitrification, estimated to account for up to ∼30% of soil N_2_O emissions under favourable conditions (Wrage et al. 2001). Nevertheless, heterotrophic denitrification is generally regarded as the major source of N_2_O in many agricultural soils, although the relative contributions of nitrification and denitrification vary with soil conditions (Weiske et al. 2001). This multi-source nature of N_2_O makes its attribution to any single functional group inherently more difficult than for NO (Butterbach-Bahl et al. 2013; Kool et al. 2011; Weiske et al. 2001). To mitigate nitrification-driven N losses, nitrification inhibitors such as 3,4-dimethylpyrazole phosphate (DMPP) and dicyandiamide (DCD) are widely applied in agricultural soils (Amanatidou et al. 2025; Zhou et al. 2020). These compounds are commercially formulated for co-application with synthetic and liquid organic fertilisers (Toda et al. 2026). Their primary mode of action involves suppression of ammonium (NH_4_□) oxidation to nitrate (NO□□) by inhibiting the activity of ammonia monooxygenase (AMO), the key enzyme initiating nitrification. This inhibition is thought to occur either through direct binding with the active site of the enzyme or by chelating the copper (Cu) cofactors required for AMO activity (Ruser and Schulz 2015). As a result, their application reduces nitrate leaching and N_2_O emissions while enhancing ammonium retention in soil systems (Subbarao et al. 2006; Zerulla et al. 2001). However, their effectiveness varies depending on soil physicochemical properties and the composition of the ammonia-oxidising community (Di and Cameron 2016). Several studies suggest that DMPP often exert stronger inhibitory effects on AOB than on AOA (Chen et al. 2015; Di and Cameron 2016; Di et al. 2010). In contrast, the effects of dicyandiamide (DCD) on AOA and AOB appear to be more variable and soil dependent(Liu et al. 2013). However, the extent to which AOB contribute to NO emissions in soils dominated by AOA remains unclear, particularly under conditions where AOB represent only a minor fraction of the nitrifying community. Moreover, because AOA and AOB differ in their physiology and sensitivity to nitrification inhibitors, the extent to which inhibitor-induced shifts in nitrifier communities influence NO and N_2_O emissions remains poorly understood. Here, we evaluated the effects of DMPP and DCD on inorganic nitrogen dynamics and the abundance of AOA and AOB and we used a dynamic flow-through DENIS system to quantify NO and N_2_O emissions from urea- and urea + DMPP-treated soil over a 28-day incubation period. Given that AOA dominated the nitrifier community of the soil studied, yet they lack the enzymatic pathways responsible for the majority of AOB-derived NO production, we hypothesised that selective inhibition of AOB by DMPP would disproportionately reduce NO emissions and increase ammonium retention relative to the uninhibited urea treatment. Furthermore, given that N_2_O emissions from fertilised soils arise from concurrent pathways, including AOA-mediated nitrification and heterotrophic denitrification, we predicted that N_2_O emissions would show a comparatively weaker and more variable response to DMPP application than NO. We therefore sought to determine whether AOB contribute disproportionately to NO emissions in a soil where AOA numerically dominate the ammonia-oxidising community.

## 2. Methods

### 2.1. Soil collection and characterisation

Soil was collected in November 2024 from a permanent grassland located at North Wyke, Devon, UK (50°46′10″ N, 3°54′05″ W) to a depth of 15 cm. The soil was classified as a clayey pelostagnogley according to the UK soil classification system (Clayden and Hollis 1984) and consisted of 44% clay, 40% silt, and 15% sand. The soil contained 0.41% total nitrogen and 11.7% organic matter and had a pH of 6.3. Plant roots and residues were removed, and the soil was subsequently passed through a 4-mm sieve. The sieved soil was stored at 4°C until further use.

### 2.2. Denitrification Incubation System (DENIS) experiment

The experiment was carried out at Rothamsted Research (North Wyke, UK) using the Denitrification Incubation System, a specialised gas-flow soil-core incubation system in which environmental conditions are tightly controlled (Cárdenas et al. 2003; Loick et al. 2016).

Soil was packed into plastic cores (4.5 cm diameter x 10 cm height) at a bulk density of 0.8 g cm^-3^. Three soil cores were placed in a stainless-steel block with equivalent holes to avoid air-spaces. Each vessel containing three soil cores represented one replicate. Each core contained approximately 127.7 g of dry soil, corresponding to approximately 383.1 g of dry soil per vessel. Prior to treatment application, the system was flushed from below with a helium–oxygen gas mixture (He:O□, 80:20) at a flow rate of 30 mL min□^1^ for six days. Following the pre-flushing phase, the flow rate was reduced to 12 mL min□^1^ and the gas stream was redirected to flow across the soil surface. This configuration was maintained for three days prior to treatment application to establish baseline gas emissions and was continued throughout the 28-day incubation period following amendment application. The DENIS gas incubation consisted of two treatments: (1) soil amended with urea and (2) soil amended with urea plus the nitrification inhibitor 3,4-dimethylpyrazole phosphate (DMPP), with three replicate vessels per treatment. Urea was applied at a rate of 100 mg N kg□^1^ dry soil, and DMPP was applied at a rate equivalent to 1.5% of the applied urea-N. The respective amendments were dissolved in 11.7 mL of water and injected into the three soil cores through ports located in the lid of each vessel to adjust the soil moisture content to 60% water-filled pore space (WFPS). The incubation cabinet was maintained at 25°C throughout the pre-flushing and incubation periods.

#### 2.2.1. Gas analyses

Gas concentrations from each vessel were automatically measured at approximately 2-h intervals throughout the experiment. Nitrous oxide (N_2_O) concentrations were quantified using a Pye Unicam 500 gas chromatograph equipped with an electron capture detector. Nitric oxide (NO) concentrations were determined using a chemiluminescence analyser (Sievers NOA280i, GE Instruments, Colorado, USA). Gas flow rates were measured throughout the experiment and recorded for each vessel. Gas fluxes were calculated from the measured gas concentrations and flow rates and normalised to the combined soil surface area of the three cores in each vessel. N_2_O and NO fluxes were expressed as mg N_2_O m□^2^ h□^1^ and mg NO m□^2^ h□^1^, respectively.

### 2.3. Destructive soil incubation experiment

A soil incubation experiment was conducted in parallel with the DENIS experiment under the same conditions of temperature, soil moisture, and bulk density to assess the effects of DCD and DMPP on soil nitrification and nitrifier microbial dynamics over a 28-day incubation period (Fig. S1b). Four treatments were included: (1) control (water only), (2) urea (urea only), (3) urea + DCD, and (4) urea + DMPP. Soil was packed into 48 plastic cores (4.5 cm diameter x 10 cm height), each containing 127.7 g of soil (oven-dry weight equivalent), corresponding to four sampling times with three replicates per treatment at each sampling time. Soil cores were prepared in the same manner as described for the DENIS experiment (Section 2.2). Urea was applied at a rate of 100 mg N kg^−1^ dry soil, while DCD and DMPP were each applied at a rate equivalent to 1.5% of the applied urea-N. The water content of all soil cores was adjusted to 60% WFPS, accounting for the volume of the treatment solutions (or deionised water for the control). Following treatment application, all soil cores were covered with aluminium foil and incubated in the dark at 25°C for 28 days. Soil moisture was maintained at 60% WFPS throughout the incubation period by replenishing water losses every two days. Soil cores were destructively sampled on days 0, 7, 14, and 28 to determine soil inorganic nitrogen (NH_4_^+^-N and TOxN concentrations) and to collect soil for DNA extraction and subsequent molecular analyses.

### 2.3.1. Soil analyses

Initial soil mineral nitrogen concentrations were determined from three randomly collected 100 g subsamples of the bulk soil prior to core packing and moisture adjustment. At each sampling time point, 5 g of soil from each sampled incubation core was extracted with 100 mL of 2 M KCl by shaking at 170 rpm for 1.5 h. The extracts were filtered through Whatman No. 2 filter paper and analysed colorimetrically using an Aquakem 250 discrete photometric analyser (Thermo Fisher Scientific, Hemel Hempstead, UK) for NH_4_^+^-N and total oxidised N (TOxN; NO□□-N + NO□□-N). At each sampling time point, TOxN and NH□□-N concentrations measured in the water-only control were subtracted from the corresponding concentrations in the treated samples to obtain net treatment-derived mineral N concentrations. Net NH_4_^+^-N and TOxN concentrations were expressed as mg N kg^-1^ dry soil.

### 2.4. Quantitative PCR (qPCR) analysis of *amoA*

DNA was extracted from soil samples using the DNeasy PowerSoil Pro Kit (Qiagen, Germany) following the manufacturer’s instructions. DNA concentration and purity were determined using a NanoDrop 2000 spectrophotometer (Thermo Fisher Scientific, Waltham, MA, USA). Extracted DNA was stored at -20°C until further analysis. The copy numbers of bacterial and archaeal *ammonia monooxygenase* (*amoA*) genes were quantified by quantitative real-time PCR (qPCR) using a CFX96 Optical Real-Time PCR Detection System (Bio-Rad Laboratories, Inc., Hercules, CA, USA) with SYBR Green I detection chemistry. DNA extracts were diluted 100-fold, and 1–10 ng of DNA was used as a template in each reaction. Primer pairs specific for bacterial and archaeal *amoA* genes were used to quantify AOB and AOA, respectively (Table 2). PCR amplifications were carried out in 10 μL reaction mixtures consisting of 5 μL of SsoAdvanced Universal SYBR® Green Supermix (Bio-Rad Laboratories, Inc., Hercules, CA, USA), 0.5 μL of each primer (10 μM), 2 µL of nuclease-free water, and 2 μL of DNA template. The thermal cycling conditions for the AOB and AOA *amoA* genes were as follows: an initial denaturation at 95°C for 3 min, followed by 46 cycles of denaturation at 95°C for 10 s, annealing at 61.5°C for AOB or 64.5°C for AOA for 40 s, and extension at 72°C for 1 min. Melt-curve analysis was performed following amplification to verify PCR product specificity, and representative amplicons were visualised by electrophoresis on a 1% (w/v) agarose gel. PCR products were gel-purified, ligated into the pGEM-T Easy vector (Promega, Madison, WI, USA), and transformed into *Escherichia coli* TOP10 (Invitrogen, Carlsbad, CA, USA). Plasmid DNA containing the target *amoA* gene was extracted and quantified using a NanoDrop 2000 spectrophotometer. The copy number of the inserted gene was calculated based on the plasmid DNA concentration, and ten-fold serial dilutions of plasmid DNA with known copy numbers were prepared in triplicate to generate standard curves for qPCR quantification. All qPCR reactions were performed in triplicate. Amplification efficiencies were 103% and 95% for AOB and AOA *amoA*, respectively, with corresponding R^2^ values of 0.99 and 0.98.

**Table 1.** Physiochemical properties of the soil used in this study. Total C: total carbon; Total N: total nitrogen; NH_4_^+^-N: ammonium-nitrogen; NO_3_^-^-N: nitrate-nitrogen

| pH | Total C<br>(g kg<sup>-1</sup>) | Total N<br>(g kg<sup>-1</sup>) | NH<sub>4</sub><sup>+</sup>-N<br>(mg kg<sup>-1</sup>) | NO<sub>3</sub><sup>-</sup>-N<br>(mg kg<sup>-1</sup>) | Clay (%) | Silt (%) | Sand (%) |
| --- | --- | --- | --- | --- | --- | --- | --- |
| 6.3 | 35.1 | 4.1 | 13.24 | 36.9 | 44 | 40 | 15 |

**Table 2.**
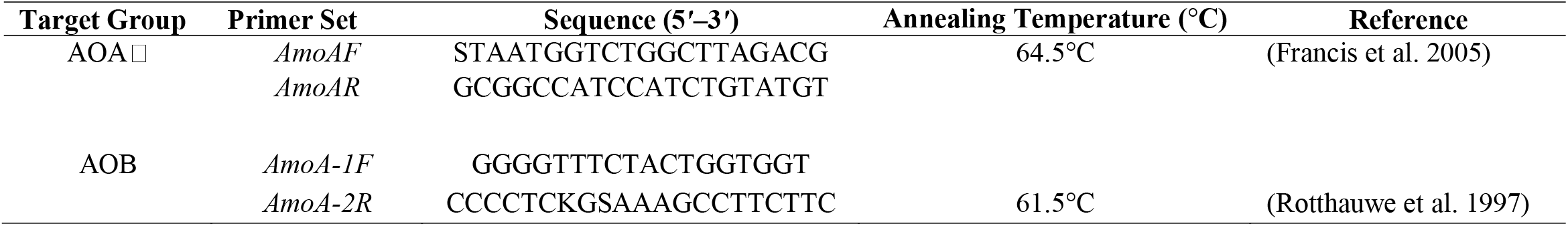
Primer sets and PCR profiles used in the real-time PCR.

### 2.5. Calculations and statistical analysis

Gas fluxes were calculated from the measured gas concentrations and flow rates and expressed as mg N_2_O m□^2^ h□^1^ for N_2_O and mg NO m□^2^ h□^1^ for NO. Flux measurements were averaged over each 24-h period to obtain daily mean fluxes for each replicate vessel. Daily treatment means were subsequently calculated from the three replicate vessels (n = 3). The percentage inhibition of N_2_O and NO fluxes by DMPP was calculated from the daily mean treatment fluxes as [(C − T)/C] × 100, where *C* is the daily mean flux of the urea treatment and *T* is the daily mean flux of the urea + DMPP treatment. NH□□ disappearance rate was calculated as [(NH□□-N at day 0) − (NH□□-N at day t)]/t, and apparent nitrification rate was calculated as [(TOxN at day t) − (TOxN at day 0)]/t, where *t* is the number of incubation days; rates were expressed as mg N kg□^1^ day□^1^. The percentage reduction in each rate by DCD or DMPP relative to urea was calculated as [(Rurea − Rinhibitor)/Rurea] × 100, where *R* is the corresponding NH□□ disappearance or nitrification rate. AOA and AOB *amoA* gene abundances, along with NH□□-N and TOxN concentrations, were tested for normality using the Shapiro–Wilk test. At each sampling time, treatment effects were analysed separately using one-way analysis of variance (ANOVA), followed by Tukey’s HSD post hoc test for multiple comparisons. Pearson correlation analysis was performed to assess the relationships among AOA and AOB *amoA* gene abundances, soil NH□□-N concentrations, and TOxN concentrations. All statistical analyses and data visualisations were conducted using GraphPad Prism software (version 10.6.0, build 796; GraphPad Software, San Diego, CA, USA). Differences were considered statistically significant at *P* < 0.05.

## 3. Results

### 3.1. Soil ammonium and total oxidised nitrogen dynamics

Dynamic changes in inorganic nitrogen forms (NH□□-N and TOxN) were observed throughout the 28-day incubation period across all treatments (Fig. 1). NH□□-N concentrations declined progressively over time in all treatments (Fig. 1a). In the urea treatment, NH□□-N decreased rapidly from approximately 75 mg kg□^1^ dry soil at day 0 to around 35, 22, and 11 mg kg□^1^ dry soil on days 7, 14, and 28, respectively. Both nitrification inhibitors significantly slowed NH□□-N depletion, resulting in higher NH□□-N concentrations than in the urea treatment at all subsequent sampling times (days 7, 14, and 28; *P* < 0.05). DMPP maintained higher NH□□-N concentrations than DCD throughout the incubation period after day 0, with significantly greater NH□□-N retention than DCD on days 7 and 28. In contrast, TOxN concentrations increased steadily during the incubation period (Fig. 1b). The urea treatment showed the greatest accumulation of TOxN, reaching approximately 95 mg kg□^1^ dry soil by day 28. Both DCD and DMPP significantly reduced TOxN accumulation relative to the urea treatment on days 7, 14, and 28 (*P* < 0.05). DMPP generally resulted in lower TOxN concentrations than DCD, with significant differences between the two inhibitor treatments on days 14 and 28 (*P* < 0.05). The calculated NH□□ disappearance rates further supported these observations (Supplementary Fig. S2a). Relative to the urea treatment, DCD reduced NH□□ disappearance by approximately 45, 50, and 19% on days 7, 14, and 28, respectively, whereas DMPP reduced NH□□ disappearance by approximately 67, 63, and 41% at the corresponding sampling times. Similarly, nitrification rates were lower in inhibitor-treated soils than in the urea treatment (Supplementary Fig. S2b). DCD reduced nitrification rates by approximately 40, 29, and 21% on days 7, 14, and 28, respectively, whereas DMPP reduced nitrification rates by approximately 46, 46, and 35%, respectively, indicating the greater efficacy of DMPP in suppressing nitrification.

**Fig. 1.**
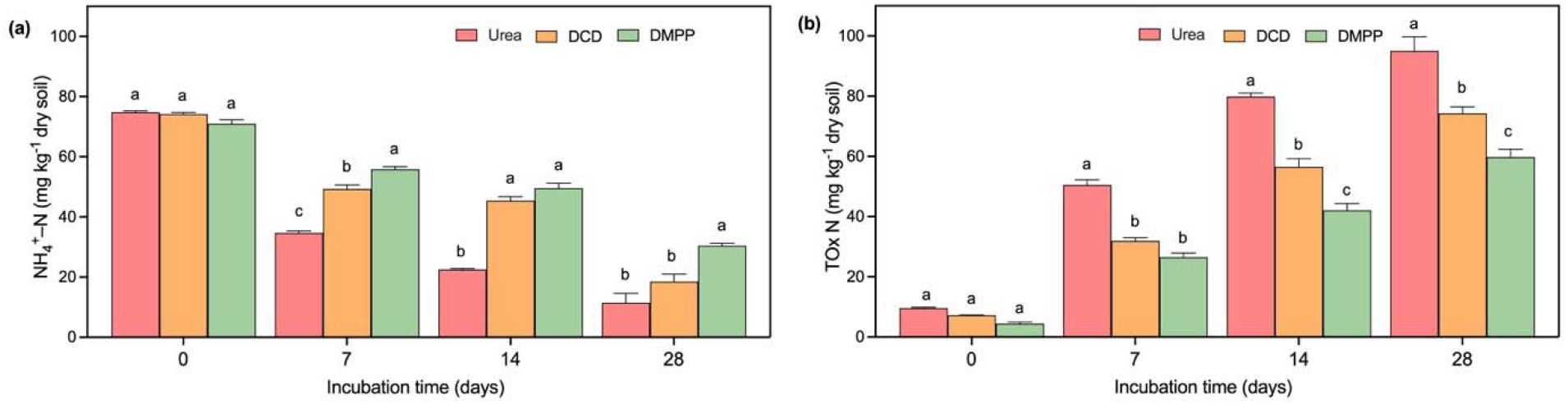
Dynamic changes in (**a**) ammonium nitrogen (NH□□-N) and (**b**) total oxidised nitrogen (TOxN) concentrations (mg kg□^1^ dry soil) under different treatments during a 28-day incubation period. Error bars represent standard deviations (n = 3) per treatment. Treatments: Urea (urea only); DCD (urea + dicyandiamide); DMPP (urea + 3,4-dimethylpyrazole phosphate). Values for fertilised treatments are expressed relative to the water control (ie. control values subtracted). Different letters above the bars within the same sampling day indicate significant differences among treatments (P < 0.05) based on one-way ANOVA and Tukey’s HSD post hoc test.

### 3.2. Abundance of ammonia oxidising archaea and bacteria

Across treatments and sampling times, ammonia-oxidising archaea (AOA) represented the larger proportion of the ammonia-oxidising community, accounting for approximately 66% based on *amoA* gene abundance, whereas ammonia-oxidising bacteria (AOB) accounted for approximately 34% (Fig. 2a). AOA *amoA* gene abundance did not differ significantly among treatments at any sampling time (Fig. 2b). In contrast, AOB *amoA* gene abundance was higher in the urea-treated soil than in the other treatments on days 7, 14, and 28 (Fig. 2c). On day 7, AOB *amoA* gene abundance was lower in the DCD- and DMPP-treated soils than in the urea-treated soil, although these differences were not significant (DCD: *P* = 0.907; DMPP: *P* = 0.227). A similar pattern was observed on day 14, with lower AOB *amoA* gene abundance in the inhibitor-treated soils, particularly in the DMPP treatment. By day 28, AOB *amoA* gene abundance was significantly lower in DMPP-treated soils than in urea-treated soils, whereas the effect of DCD was less pronounced (Fig. 2c).

**Fig. 2.**
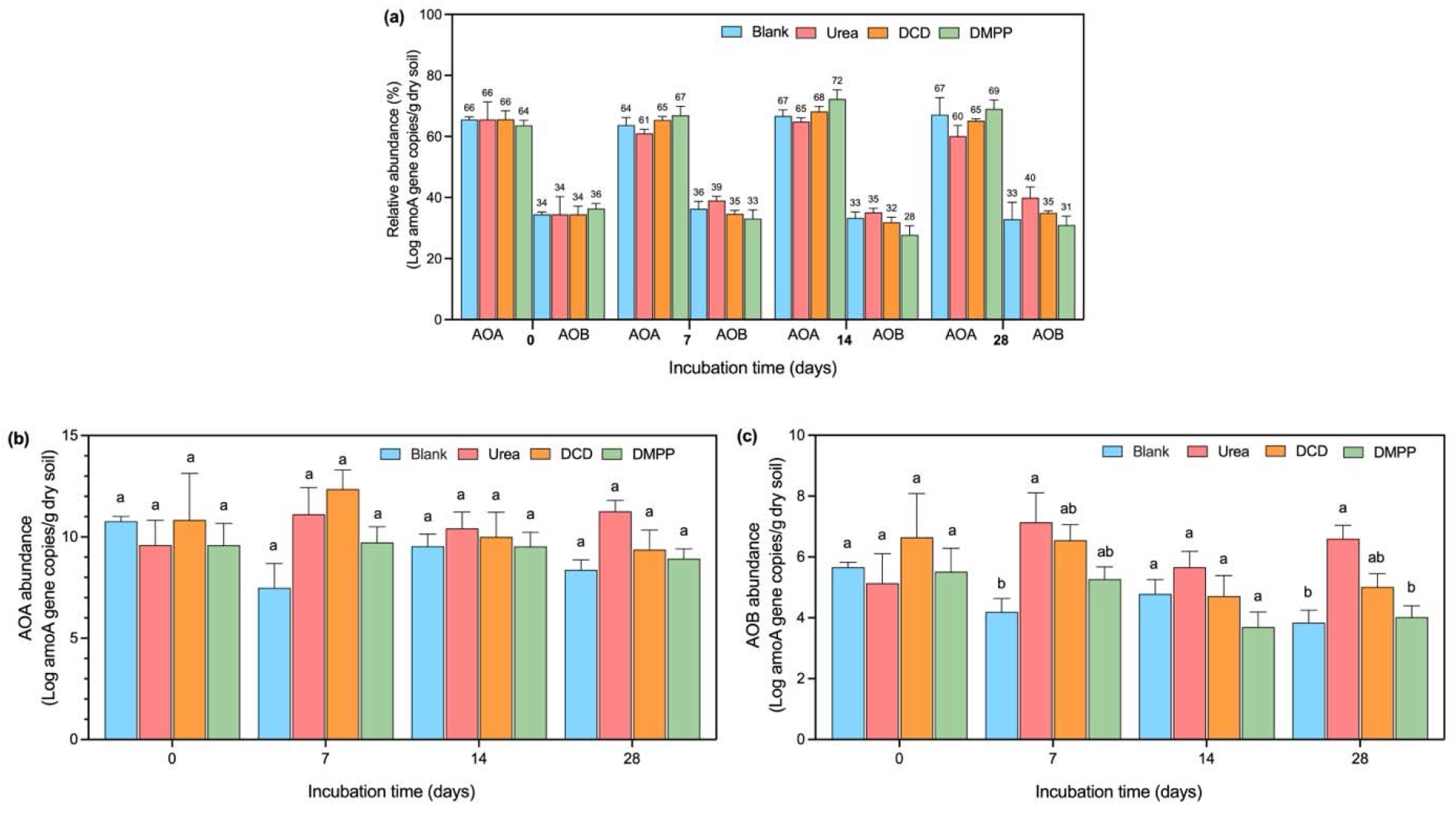
(**a**) Relative abundance (%) of ammonia-oxidising archaea (AOA) and ammonia-oxidising bacteria (AOB). (**b, c**) *amoA* gene copy abundance of AOA and AOB, respectively, under different treatments during the 28-day incubation period. Treatments were Blank (water only); Urea (urea only); DCD (urea + dicyandiamide); DMPP (urea + 3,4-dimethylpyrazole phosphate). Different letters above the bars within the same sampling day indicate significant differences among treatments (P < 0.05) based on one-way ANOVA and Tukey’s HSD post hoc test.

### 3.3. N-gas emissions

The experiment was set up to measure NO, N_2_O as well as N_2_. While both, treatment induced NO and N_2_O emissions were detected, N2 emissions remained at baseline level and are therefore not shown here.

Two treatments, urea alone and urea plus DMPP, were evaluated for gaseous N emissions using the DENIS system. N□O fluxes were consistently higher in urea-treated soil than in DMPP-treated soil throughout the 28-day incubation period (Fig. 3a). In urea-treated soil, N_2_O flux increased rapidly following fertiliser application and reached a maximum of 0.114 mg m□^2^ h□^1^ during the early stage of the incubation, followed by a gradual decline over time. In contrast, DMPP-treated soil exhibited lower N_2_O fluxes throughout the incubation period, starting at 0.083 mg m□^2^ h□^1^ and declining to 0.071 mg m□^2^ h□^1^ by day 28. Overall, DMPP reduced N_2_O flux by 0.2–27% relative to the urea treatment during the incubation period (Fig. 3c). A more pronounced response to DMPP was observed for NO (Fig. 3b). NO flux was consistently higher in urea-treated soil, reaching a maximum of 0.026 mg m□^2^ h□^1^ on day 5 before gradually declining to 0.015 mg m□^2^ h□^1^ by the end of the incubation period. In contrast, DMPP-treated soil exhibited substantially lower NO fluxes, decreasing from 0.011 mg m□^2^ h□^1^ at the start of the incubation to 0.006 mg m□^2^ h□^1^ around day 9 before gradually increasing towards the end of the incubation period. Overall, DMPP reduced NO flux by approximately 16–75% relative to the urea treatment (Fig. 3c).

**Fig. 3.**
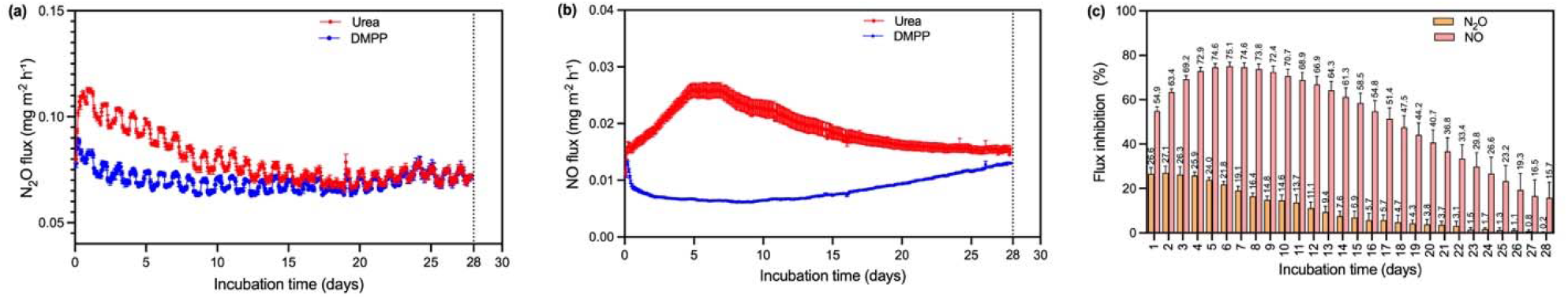
Dynamic changes in (**a)** nitrous oxide (N_2_O) flux and (**b**) and nitric oxide flux, expressed as mg m□^2^ h□^1^ from soils treated with either urea or urea plus 3,4-dimethylpyrazole phosphate (DMPP) during the 28-day incubation period. (**c**) Percentage inhibition of N_2_O and NO fluxes by DMPP relative to the urea treatment throughout the incubation period. Error bars represent ± one standard deviation (n = 3).

### 3.4. Relationships among inorganic nitrogen and ammonia-oxidiser abundances under nitrification inhibitor treatment

Pearson correlation analysis revealed distinct relationships among NH□ □-N, TOxN, and the abundances of AOA and AOB across treatments (Fig. 4). In the urea treatment (Fig. 4a), NH□ □-N was strongly and positively correlated with TOxN (r = 0.89, *P* = 0.01). Positive correlations were also observed between NH□ □-N and AOA (r = 0.45) and AOB (r = 0.65), although these were not statistically significant. Similarly, TOxN showed positive correlations with AOA (r = 0.57) and AOB (r = 0.76), while AOA and AOB abundances were positively correlated (r = 0.60). Under the DCD treatment (Fig. 4b), NH□□-N was negatively correlated with TOxN (r = −0.56), AOA (r = −0.40), and AOB (r = −0.26), although these relationships were not statistically significant. TOxN was positively correlated with AOA (r = 0.44) and AOB (r = 0.73), while AOA and AOB abundances exhibited a strong positive correlation (r = 0.84, *P* = 0.03). Under the DMPP treatment (Fig. 4c), NH□□-N was strongly negatively correlated with TOxN (r = −0.86, *P* = 0.03) and AOA abundance (r = −0.85, *P* = 0.03). A negative correlation was also observed between NH□□-N and AOB abundance (r = −0.78), although this relationship was not statistically significant (*P* = 0.06). In contrast, TOxN was positively correlated with AOB abundance (r = 0.86, *P* = 0.02), and AOA abundance was positively correlated with AOB abundance (r = 0.86, *P* = 0.02). The positive correlation between TOxN and AOA (r = 0.67) was not statistically significant.

**Fig. 4.**
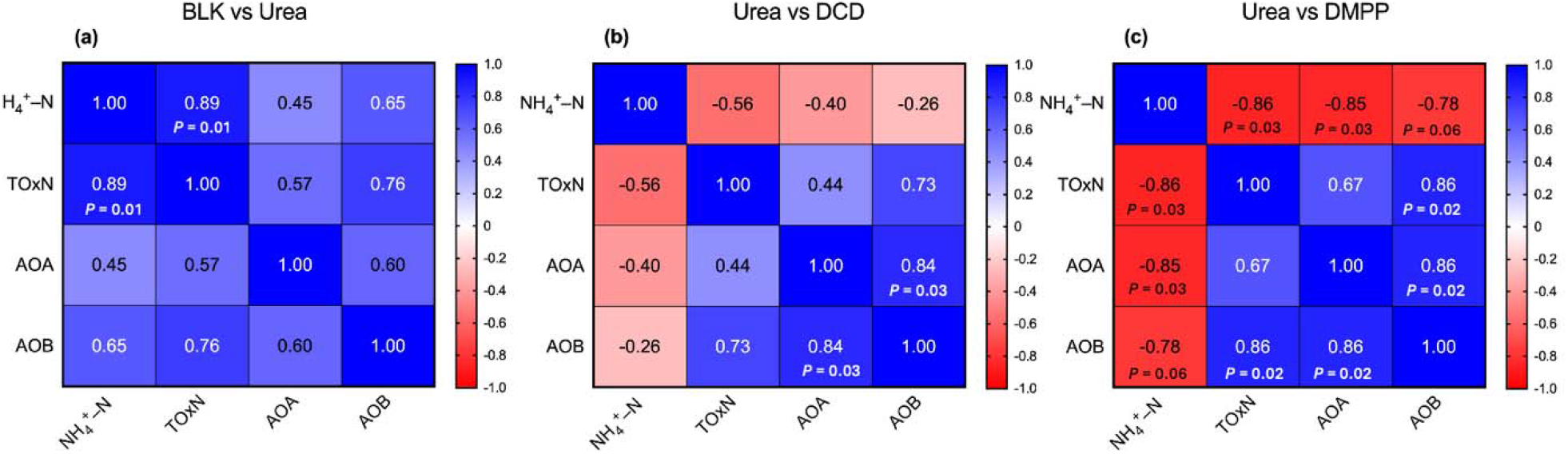
Pearson correlation matrices showing relationships among soil ammonium (NH□ □-N), total oxidise nitrogen (TOxN), and *amoA* gene abundances of ammonia-oxidising archaea (AOA) and ammonia-oxidising bacteria (AOB) at day 28. Correlations were calculated using pooled observations from (**a**) the blank and urea treatments (BLK vs Urea), (**b**) the urea and dicyandiamide treatments (Urea vs DCD), and (**c**) the urea and 3,4-dimethylpyrazole phosphate treatments (Urea vs DMPP). The colour scale indicates the strength and direction of the correlations, and the numerical values represent Pearson correlation coefficients (*r*). Statistically significant correlations (*P* < 0.05) are indicated by the corresponding *P*-values.

## 4. Discussion

### 4.1. Differential inhibition of nitrification and ammonium retention

Both DMPP and DCD suppressed nitrification over the 28-day incubation period, as evidenced by enhanced NH□□ retention and reduced TOxN accumulation relative to the urea treatment in the destructive soil incubation. However, DMPP exhibited stronger and more persistent inhibition than DCD, maintaining higher NH_4_^+^ concentrations and lower TOxN concentrations throughout the incubation period. These findings are consistent with previous studies reporting greater inhibitory efficacy and longer persistence of DMPP compared with DCD (Di and Cameron 2016; Guardia et al. 2018; Zerulla et al. 2001). The decline in NH□□-N concentrations and the corresponding accumulation of TOxN in urea-treated soil indicate rapid nitrification in the absence of nitrification inhibitors. In contrast, DMPP reduced TOxN accumulation by up to approximately 50% during the first two weeks and maintained substantial suppression until day 28, indicating prolonged inhibitory activity. DCD exhibited a weaker inhibitory effect, with an approximately 25% reduction in TOxN accumulation, possibly due to its higher mobility and faster degradation in soil (Subbarao et al. 2006; Zerulla et al. 2001). Overall, the greater inhibitory efficacy of DMPP is consistent with previous studies showing its strong inhibition of ammonium oxidation (Chen et al. 2015; Di et al. 2010; Shi et al. 2016).

### 4.2. AOA dominance and the functional importance of AOB

Although AOA constituted approximately 66% of the ammonia-oxidising community based on *amoA* gene abundance, the responses to nitrification inhibitors suggest that AOB played a functionally important role despite their lower abundance. AOA *amoA* gene abundance remained relatively stable across treatments and was not significantly affected by DMPP or DCD. These findings are consistent with previous studies showing that AOA are less sensitive to DMPP and DCD than AOB (Chen et al. 2015; Di et al. 2010). In contrast, AOB *amoA* gene abundance was lower in inhibitor-treated soils, particularly in the DMPP treatment, and this pattern coincided with reduced TOxN accumulation. This supports the concept that abundance does not necessarily equate to functional importance(Prosser and Nicol 2012). Despite their lower abundance, AOB may contribute disproportionately to nitrification under ammonium-rich conditions following urea application (Di et al. 2010). Consistent with this, AOB *amoA* gene abundance was higher in urea-treated soil than in the unfertilised control, suggesting that urea application stimulated AOB populations. Furthermore, the preferential suppression of AOB by nitrification inhibitors suggests that AOB contributed substantially to nitrification in this soil.

### 4.3. Disproportionate contribution of AOB to NO emissions

Another key finding of this study was the pronounced suppression of NO emissions by DMPP (16–75% inhibition), compared with the more moderate and variable reductions observed for N_2_O (0.2–27%). This differential response, together with the preferential effect of DMPP on AOB *amoA* gene abundance, supports our hypothesis that AOB contributed disproportionately to NO production despite their lower abundance. Nitric oxide production during nitrification is closely linked to hydroxylamine (NH_2_OH) oxidation and nitrifier denitrification (Caranto and Lancaster 2017; Stein 2011; Wrage et al. 2001). Hydroxylamine oxidation by hydroxylamine oxidoreductase can produce NO as an intermediate in AOB (Caranto and Lancaster 2017). Therefore, the preferential inhibition of AOB by DMPP likely constrained AOB-associated NO-producing pathways, contributing to the substantial reduction in NO flux observed under DMPP treatment. The sustained suppression of NO flux throughout the incubation period further suggests that AOB activity was effectively and persistently limited. These findings indicate that AOB can exert a disproportionate influence on NO production even when they are less abundant than AOA (Stein 2011). In contrast to NO, N_2_O emissions were less responsive to DMPP, exhibiting moderate inhibition (0.2–27%). This weaker response likely reflects the multiple microbial pathways contributing to N_2_O production. In soils, N_2_O can be produced via AOB-mediated nitrifier denitrification, hydroxylamine oxidation, AOA-mediated nitrification and heterotrophic denitrification (Butterbach-Bahl et al. 2013; Kool et al. 2011). Because DMPP preferentially suppresses AOB but does not strongly inhibit AOA or heterotrophic denitrifiers (Butterbach-Bahl et al. 2013; Di et al. 2010; Shi et al. 2016), alternative N_2_O-producing pathways likely remained active. This may explain why N_2_O emissions were reduced but not eliminated. The convergence of N_2_O fluxes between treatments toward the end of the incubation further suggests an increasing contribution from non-nitrifier pathways as ammonium availability declined. These findings highlight the complexity of N_2_O regulation compared with NO, supporting previous conclusions that N_2_O mitigation is more difficult due to its multiple sources(Butterbach-Bahl et al. 2013).

### 4.5. Implications for nitrogen management

Collectively, our results demonstrate that DMPP provided stronger and more sustained nitrification inhibition than DCD, leading to improved NH□□-N retention and reduced TOxN accumulation. DMPP also substantially suppressed NO emissions in the DENIS experiment. Importantly, the results suggest that AOB can contribute disproportionately to NO production despite their lower abundance, underscoring the need to consider functional activity rather than relative abundance alone when interpreting N cycling dynamics. The contrasting responses of NO and N_2_O to DMPP application further indicate that these gases respond to distinct and partially independent microbial source pools: preferential suppression of AOB may offer an effective approach for NO mitigation, whereas meaningful reductions in N_2_O will likely require integrated strategies addressing heterotrophic denitrification concurrently. From an agronomic perspective, the sustained NH□□-N retention observed under DMPP may improve nitrogen use efficiency in urea-fertilised systems, particularly in soils where AOA dominate and AOB-driven nitrification may otherwise proceed undetected under standard monitoring frameworks. Although our results indicate that DMPP suppressed NO emissions in association with reduced AOB *amoA* abundance, the specific microbial pathways responsible for NO production were not directly resolved. Future work combining ^15^N tracer approaches with isotopic source partitioning (Baggs 2008), together with measurements of *amoA* transcription and genes linked to nitrifier denitrification (e.g., *nirK* and *norB*) (Kuypers et al. 2018; Stein 2011; Wrage et al. 2001), would help to quantify the relative contributions of hydroxylamine oxidation, nitrifier denitrification, and heterotrophic denitrification to NO and N_2_O emissions. In addition, field validation across soils spanning a wider range of pH, moisture conditions, and AOA:AOB ratios will be necessary to determine how general the observed relationship between AOB and NO emissions is and to optimise inhibitor application strategies.

## 5. Conclusions

This study provides evidence that AOB, despite being less abundant than AOA, may exert a disproportionate influence on NO emissions in agricultural soil. DMPP strongly suppressed AOB abundance, reduced TOxN accumulation, enhanced NH□□ retention, and decreased NO emissions compared with urea alone. In contrast, N_2_O emissions were only moderately reduced, reflecting the contribution of multiple microbial pathways beyond AOB-mediated nitrification. These results show that microbial functional activity, rather than relative abundance, is the better predictor of gaseous N losses, and that NO and N_2_O mitigation can be decoupled depending on which pathway an inhibitor target. This points to AOB-mediated processes as a potential target for reducing NO emissions and suggests that inhibitor efficacy should be evaluated based on functional responses, including nitrification and associated gas production, rather than nitrifier abundance alone.

## Acknowledgments

The authors gratefully acknowledge SABIC for financial support of this research project (grant number RGC/3/6010-01-01). Their commitment to advancing sustainable agricultural technologies and innovative nitrogen management solutions made this work possible. We also sincerely thank Rothamsted Research, United Kingdom, for their valuable collaboration and scientific support, particularly to the EPSRC funded project: Global Nitrogen Innovation Center for Clean Energy and the Environment (NICCEE) (EP/Y025776/1). The authors also acknowledge the Analytical Chemistry Core Lab at King Abdullah University of Science and Technology (KAUST) for providing analytical support and access to instrumentation essential for this research. Finally, the authors are grateful to all colleagues and technical staff at both institutions whose assistance and support contributed to this research.

## Statements & Declarations

### 3. Author Contributions

All authors conceived the study and designed the methodology. A.A and N.L collected and processed the soil samples. A.A performed the experimental work, carried out the data analysis, and wrote the first draft of the manuscript. B.V and B.B provided financial support for the successful completion of the research. V.J.M and L.M.C contributed to study supervision, interpretation of results, and manuscript revision. All authors critically reviewed the manuscript and approved the final version for publication.

### 4. Data Availability

The datasets generated and/or analysed during the current study are available from the corresponding author on reasonable request.

## Supplementary Information

**Supplementary Figure 1.**
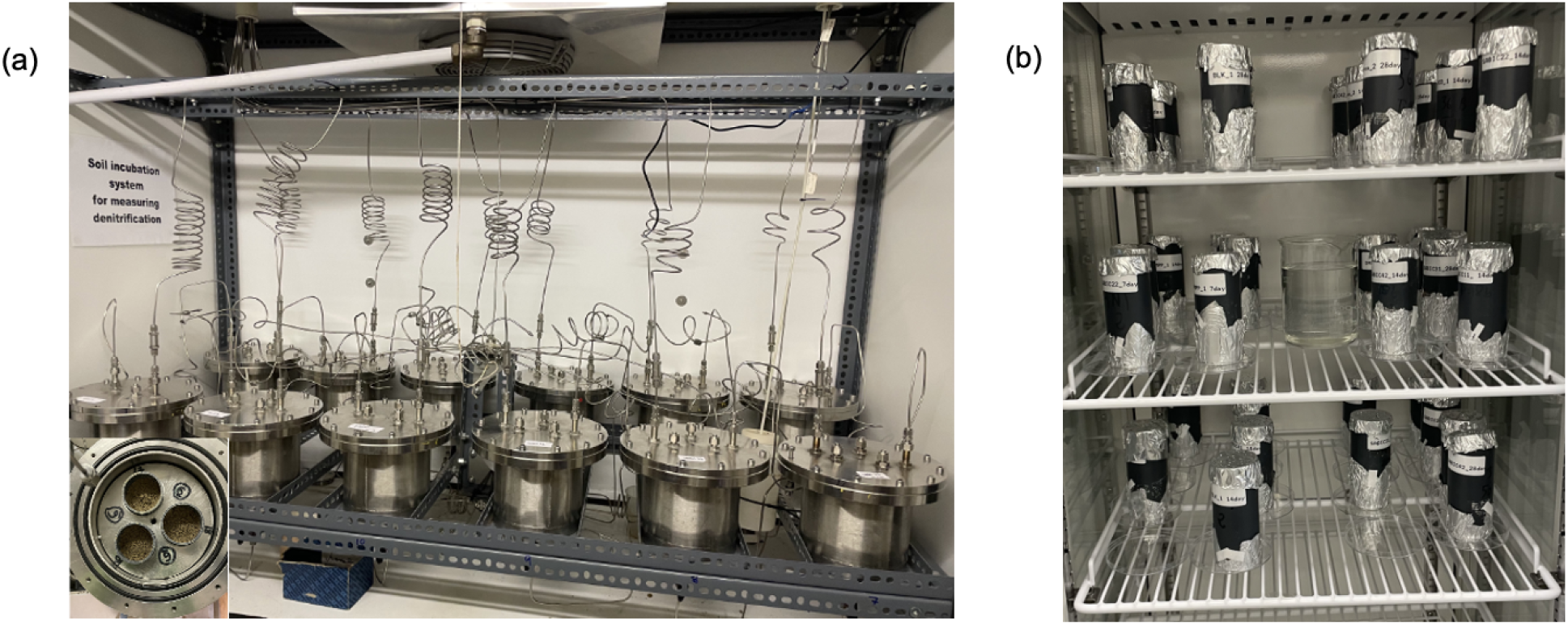
Experimental setups used in this study. (**a**) DENIS experiment showing soil cores inside the chambers. (**b**) Destructive soil incubation experiment conducted in parallel under controlled conditions (25□°C and relative humidity was maintained between 60-70%

**Supplementary Figure 2.**
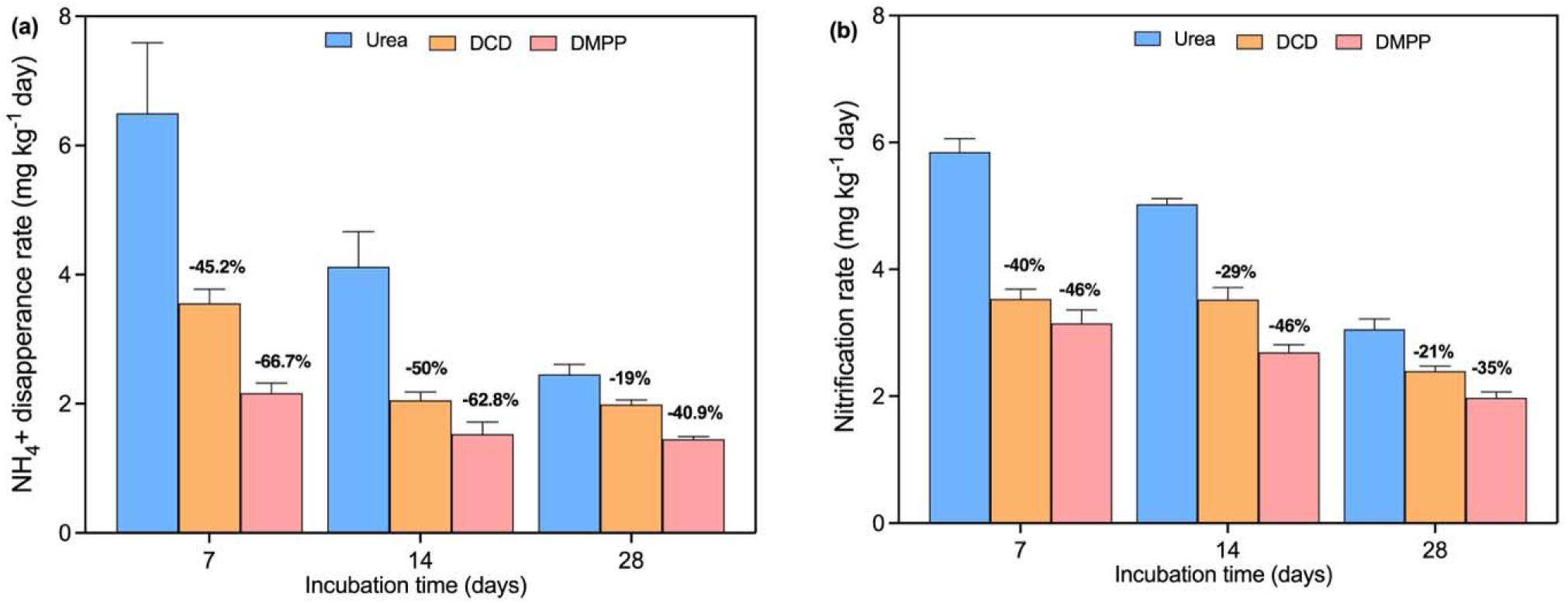
(**a**) Ammonium disappearance rate and (**b**) nitrification rate under DCD and DMPP treatments on days 7, 14, and 28. Error bars represent standard deviations of the means (n = 3). Percentages shown above the bars represent the effectiveness of DCD and DMPP in reducing ammonium disappearance and nitrification rates relative to the urea treatment.

